# Inhibition of GCN5 decreases skeletal muscle fat metabolism during high fat diet feeding

**DOI:** 10.1101/2022.10.18.512735

**Authors:** Alexander E. Green, Brayden L. Perras, Hongbo Zhang, Elena Katsyuba, Alaa Haboush, Kwadjo M. Nyarko, Dheeraj K. Pandey, Abolfazl Nik-Akhtar, Dongryeol Ryu, Johan Auwerx, Keir J. Menzies

## Abstract

**Introduction:** GCN5 (*Kat2a*) is a lysine acetyl transferase capable of acetylating and inhibiting PGC-1α activity. As such, it is described as a negative regulator of PGC-1α and subsequently restricts mitochondrial content. However, elimination of GCN5 in skeletal muscle does not increase mitochondrial content or alter lipid metabolism under normal metabolic conditions. GCN5 levels increase with high-fat diet (HFD) feeding in rodents. Additionally, the GCN5 homolog, PCAF, has previously been shown to also acetylate and inhibit PGC-1α and therefore may possibly compensate for loss of GCN5.

**Objective:** The objective of this study was to examine if with HFD feeding that elimination of GCN5 (*Kat2a* gene) from skeletal muscle would elicit improvements in mitochondrial and metabolic markers.

**Methods:** Skeletal muscle specific GCN5 knockouts (*Gcn5 skm-/-*) were fed an HFD. Body composition, cardio-metabolic and physical fitness outcomes were monitored. Additionally, cultured myotubes were treated with a pan-GCN5/PCAF inhibitor and examined for changes in mitochondrial markers.

**Results:** Elimination of skeletal muscle GCN5 did not alter body composition, tissue masses, energy intake, or energy expenditure measurements of mice fed an HFD. Furthermore, whole body glucose homeostasis and cardiac measurements were not altered. There were few differences in lipid metabolism genes, relatively more glucose oxidation versus *Gcn5 skm+/+* (wildtype) mice, and a reduction in *Pdk4* expression. Exercise capacity and mitochondrial content levels were not altered in *Gcn5 skm-/-* mice. Further, elimination of GCN5 in skeletal muscle increased *Kat2b* (PCAF) mRNA expression; however, inhibition of GCN5/PCAF bromodomains in cultured myotubes did not increase oxidative metabolism genes and decreased expression of some mitochondrial genes and *Pdk4* mRNA.

**Conclusions:** Neither elimination of GCN5, nor simultaneous inhibition of GCN5 and its homolog PCAF improved skeletal muscle mitochondrial content under normal or HFD-fed conditions. Despite this, GCN5 may play a role in regulating macronutrient preference by regulating *Pdk4* content. Thus, HFD/macronutrient excess revealed novel roles of GCN5 in skeletal muscle.

**Highlights:**

– Skeletal muscle specific elimination of *Gcn5/Kat2a* decreases fat oxidation without 1) preventing high-fat diet induced weight gain, 2) improving whole body glucose handling, or 3) improving skeletal muscle mitochondrial content.
– Inhibition of the GCN5 and PCAF bromodomains and *Gcn5* ablation decreases expression of *Pdk4*.
– Expression of *Kat2b* increases with *Gcn5* elimination in skeletal muscle.
– Inhibition of the GCN5 and PCAF bromodomains do not result in increased skeletal muscle mitochondrial content.

## 1 Introduction

Protein acetylation is a highly conserved post-translational modification that is intricately intertwined with cellular metabolism[1]. The acetylation status of proteins is governed by a balance between the activity of protein acetyltransferases and deacetylases, as well as the availability of their substrates including acetyl-CoA and Nicotinamide adenine dinucleotide (NAD+)[2]. Further, acetylation and deacetylation of certain proteins, such as PGC-1α – a major transcriptional coordinator of cellular metabolism and mitochondrial content – are thought to regulate cellular metabolism and availability of the substrates for acetyltransferases/deacetylases. This further illustrates the interconnected path of acetylation and metabolism. Despite extensive interest and research into how acetylation controls key metabolic proteins, identification of single regulatory acetyltransferases/deacetylases for therapeutic targeting and consequently development of effective inhibitors/activators to alter cellular metabolism have proven thus far elusive.

Previously, gain of function models revealed GCN5 as a major regulator of cellular metabolism via acetylation and inhibition of PGC-1α activity[3–5]. However, recent studies revealed that under basal chow-fed conditions, elimination of GCN5 alone[6], as well as in combination with overexpression of *Sirt1* (a PGC-1α deacetylase)[7] had relatively minor effects on overall metabolism. As GCN5 expression has previously been described to increase with high fat diet (HFD)-feeding[8] and some HFD-feeding regimens increase mitochondrial content in muscle[9], herein skeletal muscle specific GCN5 ablated (*Gcn5 skm-/-*) mice[10] were challenged with HFD-feeding to investigate if a HFD exacerbated or revealed differences in cellular/energy metabolism. Further, due to the high homology between GCN5 and PCAF[11], we examined the effect of dual inhibition of both proteins on mitochondrial content. We hypothesized 1) that under an HFD, *Gcn5 skm-/-* mice would have superior fat oxidation and mitochondrial content, and subsequently improved cardio-metabolic parameters; and 2) that PCAF may compensate for the lack of GCN5 in skeletal muscle and thus, dual-inhibition of GCN5 and PCAF would reveal a greater improvement in skeletal muscle mitochondrial content.

## 2 Materials and Methods

### 2.1 Animal Housing

All animal experiments were approved by the University of Ottawa Animal Care Committee (Protocols #: Hse-3236 and Hse-3607) and the ethics committee of the Canton de Vaud, Switzerland (Permit # 2285). Generation of *Gcn5 skm-/-* mice has been previously described[10] and used *Gcn5* floxed mice (Jackson Laboratory: 033454) and HSA-Cre mice (Jackson Laboratory: 006149). Mice were genotyped using the primers included in Table S3 Herein *Gcn5* floxed, HSA-Cre mice are used as control mice and referred to as *Gcn5 skm+/+. Gcn5* floxed, HSA-Cre^+/-^ are referred to as *Gcn5 skm-/-*. Animals were housed at the University of Ottawa or the École Polytechnique Fédérale de Lausanne on a 12-hour/12-hour light/dark cycle at ~23°C with *ad libitum* water and food access.

Mice were fed a control diet (Research Diets D12450J) or high fat diet (60% of kcal from fat; Research Diets D12492) as indicated. All animal phenotyping experiments were completed according to the standard operating procedures of the Eumorphia program (https://www.eumorphia.org) [12]. Tissues were collected from mice in the fed state between 9 am and noon.

### 2.2 Cell Culture Experiments

Primary myoblasts were purified from C57Bl/6NTac mice using a pre-plating technique. Briefly, muscles were dissected from the hindlimb and thoroughly minced. Subsequently, the muscles were digested in a solution of 10 g/L Collagenase B (11088831001 Roche Canada, Basel, Switzerland) and 4 g/L Dispase II (D4693 Millipore Sigma, Burlington, Massachusetts, USA) in Ham’s F10 Media (318-050-CL, Wisent, Saint-Jean Baptiste, QC, Canada) at 37°C for ~30 minutes with occasional trituration. Digested solution was passed through a 100 μm cell strainer and resuspended in growth media consisting of Ham’s F10 Media supplemented with 20% Bovine calf serum (SH3007203, Marlborough, Massachusetts, United States), 1% penicillin-streptomycin and 2.5 ng/mL basic Fibroblast Growth Factor (ab155734, Abcam, Cambridge UK). The cell suspension was plated for 1-2 hours on a non-collagen coated plate. The supernatant that contained an enriched fraction of myoblasts was transferred to a collagen coated plate for 24 hours. Subsequently, cells were removed using 0.05% trypsin-EDTA solution (325-043-CL, Wisent Inc., Saint-Jean-Baptiste, QC) and underwent a second pre-plating for 1-2 hours on non-collagen coated plates. The supernatant was then transferred to collagen coated plates and used for subsequent experiments.

For murine primary myoblast derived-myotube (herein primary myotubes) experiments, cells were plated at ~100,000 cells/cm^2^ in growth media and allowed to adhere for ~16 hours. Cells were then incubated in differentiation media consisting of high glucose Dulbecco’s Modification of Eagle Medium (319-005-CL, Wisent, Saint-Jean Baptiste, QC, Canada) supplemented with 5% heat inactivated horse serum (26050088, Gibco™ Fisher Scientific Company, Ottawa, ON, Canada) and 1% penicillin=streptomycin. Differentiation media was refreshed daily for 3 days. After 72 hours, cells were treated with either GSK4028 (HY-101027A, Med Chem Express, Monmouth Junction, NJ, USA) or GSK4027 (HY-101027, Med Chem Express, Monmouth Junction, NJ, USA)[13] at the indicated doses for 72 hours prior to harvest as described below.

### 2.3 Tissue Histology

Approximately 50 mg chunks of gonadal adipose tissue and liver tissue were excised from animals and submerged in 10% formalin for > 24 hours. Subsequently, tissues were moved to 70% EtOH and sent to the Louis Pelletier Histology Core for paraffin embedding, 4 μm sectioning, and hematoxylin and eosin staining. Images were obtained using a Axio Scan Z1 slide scanner at 20x (Zeiss, Oberkochen, Germany). FIJI was used to quantify the size of white area within the liver sections as a marker of lipid deposition. For adipose tissue, 250 adipocytes were quantified for minimum feret diameter at random by a blinded individual using FIJI.

### 2.4 Protein Quantification

For tissue analysis, flash frozen quadriceps muscle was first pulverized on a liquid nitrogen cooled aluminum block. Approximately 15-25 mg of pulverized tissue was weighed into a reinforced homogenization tube with 2 ceramic beads. Immediately before homogenization, radioimmunoprecipitation (RIPA) buffer supplemented with Roche cOmplete Protease Inhibitor Cocktail (05892970001, Roche Canada, Basel, Switzerland), Roche PHOStop Phosphatase Inhibitor Cocktail (4906837001 Roche Canada, Basel, Switzerland), and 5 mM Sodium Butyrate at a volume of 10 μL/mg of tissue. Homogenization tubes were shaken on a Fisher Scientific Bead Mill 24 (Fisher Scientific Company, Ottawa, ON, Canada). For myotubes protein extraction, cells were rinsed twice with PBS and RIPA immediately applied. Cells were flash frozen and frozen lysate scraped and transferred to 1.5 mL centrifuge tubes. Following homogenization/scraping, tubes were immediately placed on ice and then centrifuged at 16,000 x g at 4°C and the supernatant was removed. This was performed twice.

Total protein was quantified using the DC protein assay (Bio-Rad Laboratories, Hercules, California, USA). Subsequently, protein lysates were prepared for western blotting by diluting samples in 4x Laemmli Buffer (Bio-Rad Laboratories, Hercules, California, USA), RIPA and β-mercaptoethanol. Diluted lysates were then separated by size on FastCast SDS-PAGE gels (Bio-Rad Laboratories, Hercules, California, USA). Separated proteins were transferred to nitrocellulose membranes using a TurboTransfer apparatus (Bio-Rad Laboratories, Hercules, California, USA). Non-specific binding of proteins was blocked for 1 hour at room temperature using blocking buffer consisting of 5% bovine serum albumin (BSA, AD0023, BioBasic, Markham, ON, Canada) diluted in Tris-Buffered-Saline with 0.1% Tween-20 (TBS-T). Membranes were interrogated with primary antibodies (Table S5) diluted in 5% BSA diluted in TBST-T overnight at 4°C. Membranes were then washed with TBS-T and incubated with secondary antibodies conjugated to horse radish peroxidase for 1 hour at room temperature. Detection was performed on a ChemiDoc system (Bio-Rad Laboratories, Hercules, California, USA) using Clarity (Bio-Rad Laboratories, Hercules, California, USA) or Clarity Max (Bio-Rad Laboratories, Hercules, California, USA) enhanced chemiluminescent solutions. Quantification of blots was performed using FIJI following rolling ball background subtraction.

### 2.5 RNA Analysis

For tissue analysis, quadriceps muscle was weighed out into homogenization tubes as described above. Prior to homogenization, 1 mL of TRIzol™ (15596018, Fisher Scientific Company, Ottawa, ON, Canada) was added to each tube. 200 μL of chloroform was added to each tube and the tubes were inverted 10 times. Samples were centrifuged at 4°C and 16,000 x g for 15 min. The aqueous phase was removed and further purified using an EZ-10 DNAaway RNA Mini Prep Kit (BS881133, BioBasic, Markham, ON, Canada) as per the manufacturer’s instructions. For RNA extraction from myotubes, was extracted only using an EZ-10 DNAaway RNA Mini Prep Kit (BS881133, BioBasic, Markham, ON, Canada). RNA was eluted in RNAase-free water and quantified using a NanoDrop 2000 Spectrophotometer (Fisher Scientific Company, Ottawa, ON, Canada).

cDNA synthesis was completed using a NEB First Strand cDNA synthesis kit (M0368; New England Biolabs Canada, Whitby, ON, Canada). Briefly, 2 ug of mRNA was aliquoted and incubated with ProtoScript Reverse Transcriptase, 62.5 μM Random Hexamers, 0.5 mM dNTPs, 10 mM DTT, and ProtoScript II Buffer at 25°C for 5 minutes, 42°C for 60 minutes and 65°C for 20 minutes.

For Sybr Green-based qPCR, Luna Taq Premade Master Mix was used (M3003E, New England Biolabs Canada, Whitby, ON, Canada). Briefly, 2.5 μL of Luna Taq Premade Master Mix, 0.25 μM of Forward and Reverse primers, and 2 μL of RNAse/DNAse-free water was combined with a 2.5 ng of cDNA. Primer sequences were obtained from PrimerBank (https://pga.mgh.harvard.edu/primerbank/)[14] and confirmed using PrimerBlast (https://www.ncbi.nlm.nih.gov/tools/primer-blast/) for specificity and where possible exon-spanning (Table S1).

For Taqman probe-based qPCR, a master mix containing 0.25 U Amplitaq Gold (4486226 Thermo Fisher Scientific, Waltham, Massachusetts, United States), 2 μmol of dNTPs, 25 μmol of MgCl2, and a 0.25 dilution of Taqman Probe (Table S2) was made and combined with 2.5 ng of cDNA.

All reactions were performed and quantified on a CFX386 QPCR System (Bio-Rad Laboratories, Hercules, California, USA) with the following cycling parameters: 95°C for 1 minute, then 45 cycles of: 95°C for 15 seconds and 60°C for 30 seconds.

### 2.6 mtDNA and nDNA Quantification

Total genomic DNA was extracted with an EZ-10 Spin Column Animal Genomic DNA Miniprep Kit (BS427, BioBasic, Markham, ON, Canada) as per the manufacturer’s instructions. Quantification of mtDNA to nDNA ratio was performed as previously described [15]. Primer sequences are included in Table S4. The same master mix and cycling parameters were used as in the RNA analysis, except that 5 ng of gDNA was used as a template.

### 2.7 Statistical Analysis

All statistical analysis was performed using GraphPad Prism 9. For comparisons between 2-groups at one time point, student’s t-tests were used. For comparisons between 2 groups over multiple measurement points, 2-way repeated measures analysis of variances (ANOVA) were performed. For comparisons between 2 groups and 2 conditions (i.e., diet and genotype) at a single time point, 2-way ANOVAs were completed. For all ANOVAs, Bonferroni post hoc tests comparing differences only between genotypes were completed unless otherwise indicated. For every test, an α value of 0.05 was used to determine statistical significance. Any comparisons with p-values < 0.1 are indicated on graphs and reported as not-statistically significant trends.

## 3 Results

HFD-feeding has previously been described to upregulate the mRNA and protein expression of GCN5[8]. Since GCN5 levels are elevated under a HFD and previous reports have found no metabolic differences in muscle specific GCN5 KOs fed chow diets[6,7], in this study *Gcn5 skm-/-* mice were metabolically challenged with HFD-feeding.

Consistent with previous reports[10], *Gcn5/Kat2a* mRNA levels, assessed with multiple primer sets, were reduced in quadriceps muscle of *Gcn5 skm-/-* (Fig. 1A and Fig. S1A) in both control- and HFD-fed mice. However, there was a trend for an increase in *Gcn5* levels with HFD-feeding that, in contrast to previous studies[8], did not reach statistical significance. No differences between genotypes were observed in body mass (Fig. 1B), lean mass (Fig. 1C), fat mass (Fig. 1D) or tissue masses (Fig. 1E). Additionally, adipocyte size (Fig. 1F-G) and liver white area (a surrogate marker of liver lipid content; Fig. 1H-I) were not different between genotypes. Consistent with no differences in body composition, there were no differences in oxygen consumption rates (Fig. 1J) or physical activity levels (Fig. 1K). Interestingly, there was an overall decrease in oxygen consumption rates when corrected for activity levels, suggesting decreased basal metabolism (Fig. 1L); however, this difference was small enough to not affect overall body mass (Fig. 1B).

**Figure 1.**
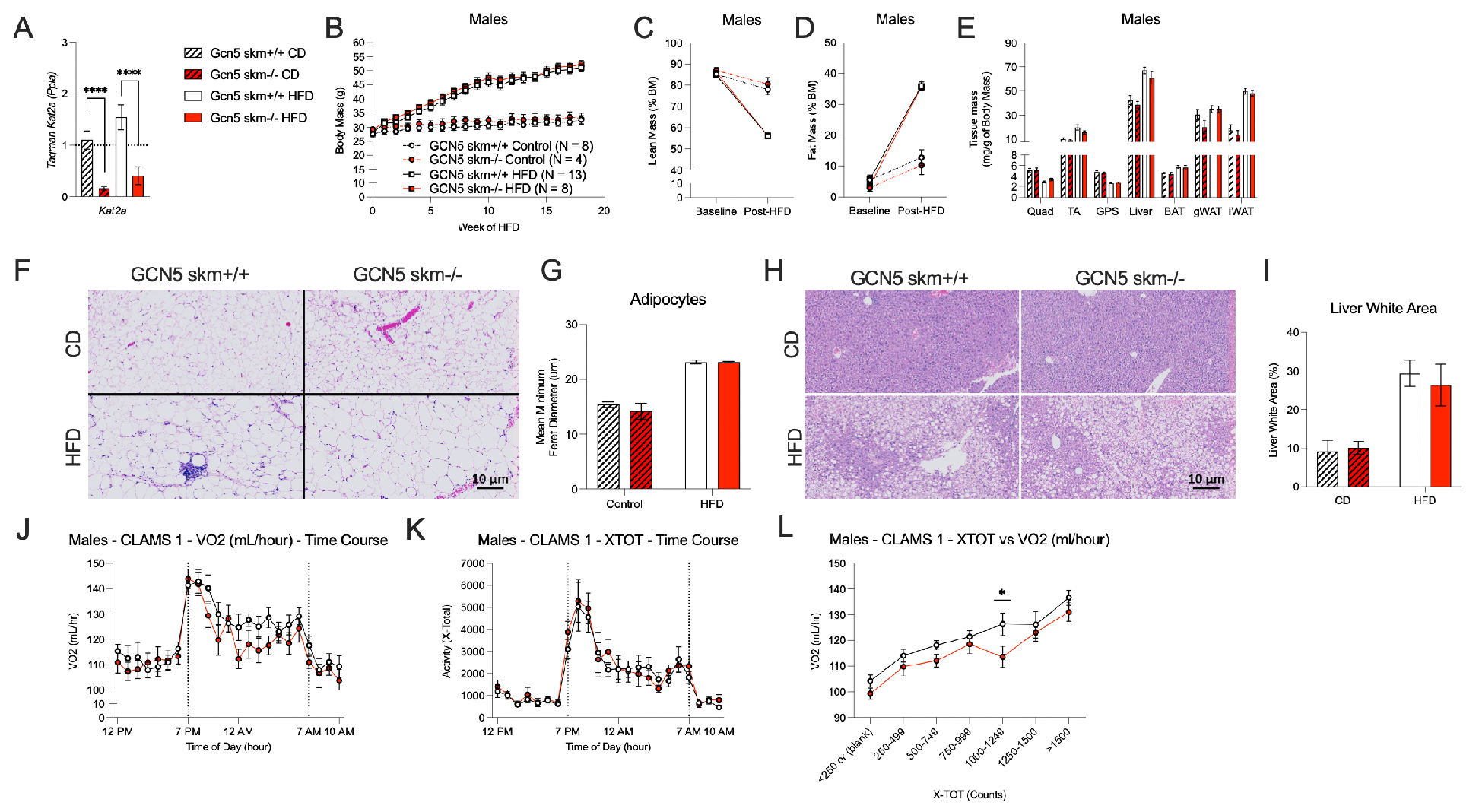
Mice lacking GCN5 in skeletal muscle gain similar lipid mass and expend similar levels of energy at all activity levels. A) Kat2a mRNA is decreased in quadriceps muscle of Gcn5 skm-/- mice (n = 4-13). Body Mass (B), lean mass (C), fat mass (D) and all tissue masses (E) were not significantly different between male Gcn5 skm+/+ and Gcn5 skm-/- mice (n = 4-13). Adipocyte size (F and G) and liver (H and I) white area were not different between genotypes (n = 3). No genotype differences were observed in HFD-fed mice in oxygen consumption rates (J) or total physical activity (K) but an overall decrease in oxygen consumption independent of activity (L; n = 12). Bars in graphs represent mean values and error bars are s.e.m. *: p < 0.05, ** p < 0.01, *** p < 0.001 and **** p < 0.0001 indicates significance by 1/2-way ANOVA and Bonferroni post-hoc tests.

To investigate the origin of this small reduction in basal metabolism, respiratory exchange ratio (RER) was measured. As there was a slight increase in RER (Fig. 2A), this suggested there was a greater preference for carbohydrate oxidation in *Gcn5 skm-/-* mice than *Gcn5 skm+/+*. Additionally, expression of *Pcx* (a gene supporting glucose derived TCA-cycle anaplerosis) and *Pdk4* (a gene inhibiting pyruvate entry into the TCA-cycle) were reduced and this may consequently permit increased glucose oxidation. Further, genes supporting *de novo* lipogenesis and glycerolipid synthesis (i.e., *Acacb, Acly*, and *Gpam*) were particularly reduced in *Gcn5 skm-/-* mice (Fig 2B). Together, these findings suggest that *Gcn5 skm-/-* mice have a relatively greater preference for carbohydrate oxidation versus *Gcn5 skm+/+* mice and loss of Gcn5 may prevent entry of TCA-cycle derived metabolite into lipogenesis.

**Figure 2.**
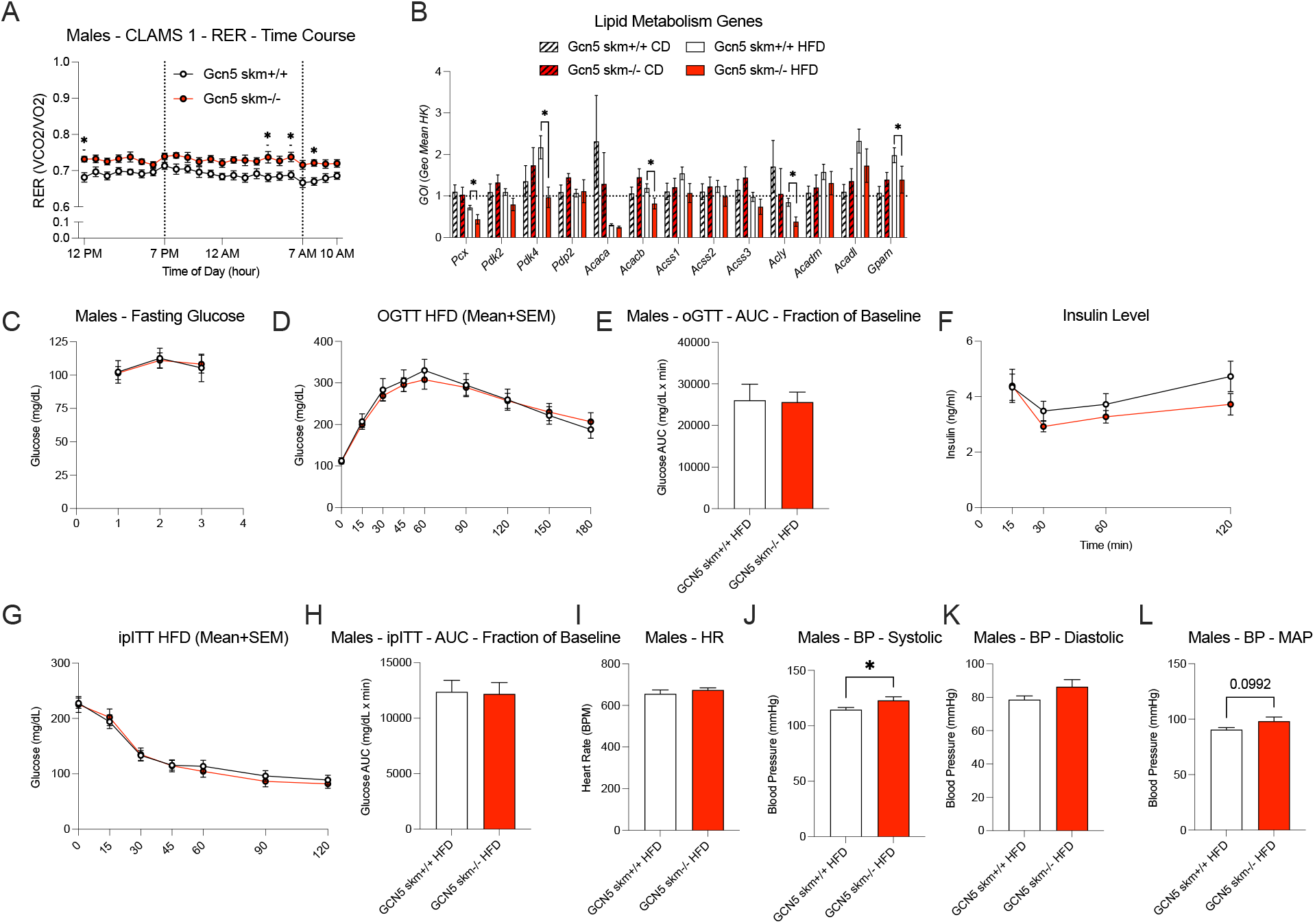
GCN5 ablation in skeletal muscle reduces fat oxidation/increases glucose oxidation but does not significantly affect glucose handling, glucose tolerance, or cardiovascular parameters. A) Gcn5 skm-/- mice have elevated RER (n = 12). Quadriceps mRNA measures (n = 4-12) of lipid metabolism genes (B). No differences were observed between Gcn5 skm+/+ and skm-/- mice in fasting blood glucose (C), glucose tolerance (D-E), glucose stimulated insulin secretion (F), and insulin tolerance (G-H; all n = 12). No difference between genotypes in for heart rate (I), diastolic blood pressure (K) and mean arterial pressure (L), but there was a significance difference in systolic blood pressure (J; all n = 12). Bars in graphs represent mean values and error bars are s.e.m. *: p < 0.05, ** p < 0.01, *** p < 0.001 and **** p < 0.0001 indicates significance by 1/2-way ANOVA and Bonferroni post-hoc tests.

Despite the absence of changes in overall body mass, *Gcn5 skm-/-* mice were examined for changes in cardio-metabolic parameters that can be altered independent of body mass. Although *Gcn5 skm-/-* mice had reduced levels of *Pcx* and *Pdk4*, there were no differences in fasted blood glucose levels (Fig. 2C) or glucose tolerance (Fig. 2D-E). Further, glucose-stimulated insulin secretion (Fig. 2F) and insulin sensitivity (Fig. 2G-H) were not improved in *Gcn5 skm-/-* mice. Additionally, heart rate (Fig. 2I) and diastolic blood pressure (Fig. 2K) were not altered; however, systolic blood pressure (Fig. 2J) and mean arterial pressure (Fig. 2L) were increased or had a trend for an increase.

As GCN5-mediated acetylation has specifically been described to inhibit PGC-1α activity[5,16] and skeletal muscle endurance capacity is highly dependent upon mitochondrial content, *Gcn5 skm-/-* mice were assessed for their running capacity and mitochondrial content following HFD-feeding. Despite the slightly diminished lipid oxidation rates, maximal running tests to exhaustion did not reveal any improvements in time to exhaustion or distance at exhaustion in *Gcn5 skm-/-* mice (Fig. 3A and B), consistent with previous reports in chow-fed *Gcn5* muscle KO mice[6,7]. Additionally, as acute thermogenesis is dependent upon skeletal muscle contractile activity when shivering, *Gcn5 skm-/-* were housed at 4°C and monitored for decreases in rectal temperature as another test of skeletal muscle function. No differences were observed between *Gcn5 skm+/+* and *Gcn5 skm-/-* mice (Fig. 3C). To further evaluate if, at the molecular level, there were differences in genes known to be regulated by GCN5, levels of transcription factors directly and indirectly regulated by acetylation were measured. In agreement with decreased amounts of fat oxidation and *de novo lipogenesis* genes, levels of *Ppargc1a, Sirt1, Srebf1* and *Nfatc3* were decreased in HFD-fed *Gcn5 skm-/-* mice versus *Gcn5 skm+/+* mice (Fig. 3D). However, there were no differences in representative mitochondrial oxidative phosphorylation protein levels (Fig. 3E), mtDNA levels (Fig. 3F) or structural mitochondrial genes (Fig. 3H). Though, we observed decreased levels of certain genes involved in oxidative phosphorylation (Fig. 3G). Altogether, these results suggest that overall mitochondrial content was not different or slightly reduced in *Gcn5 skm-/-* mice.

**Figure 3.**
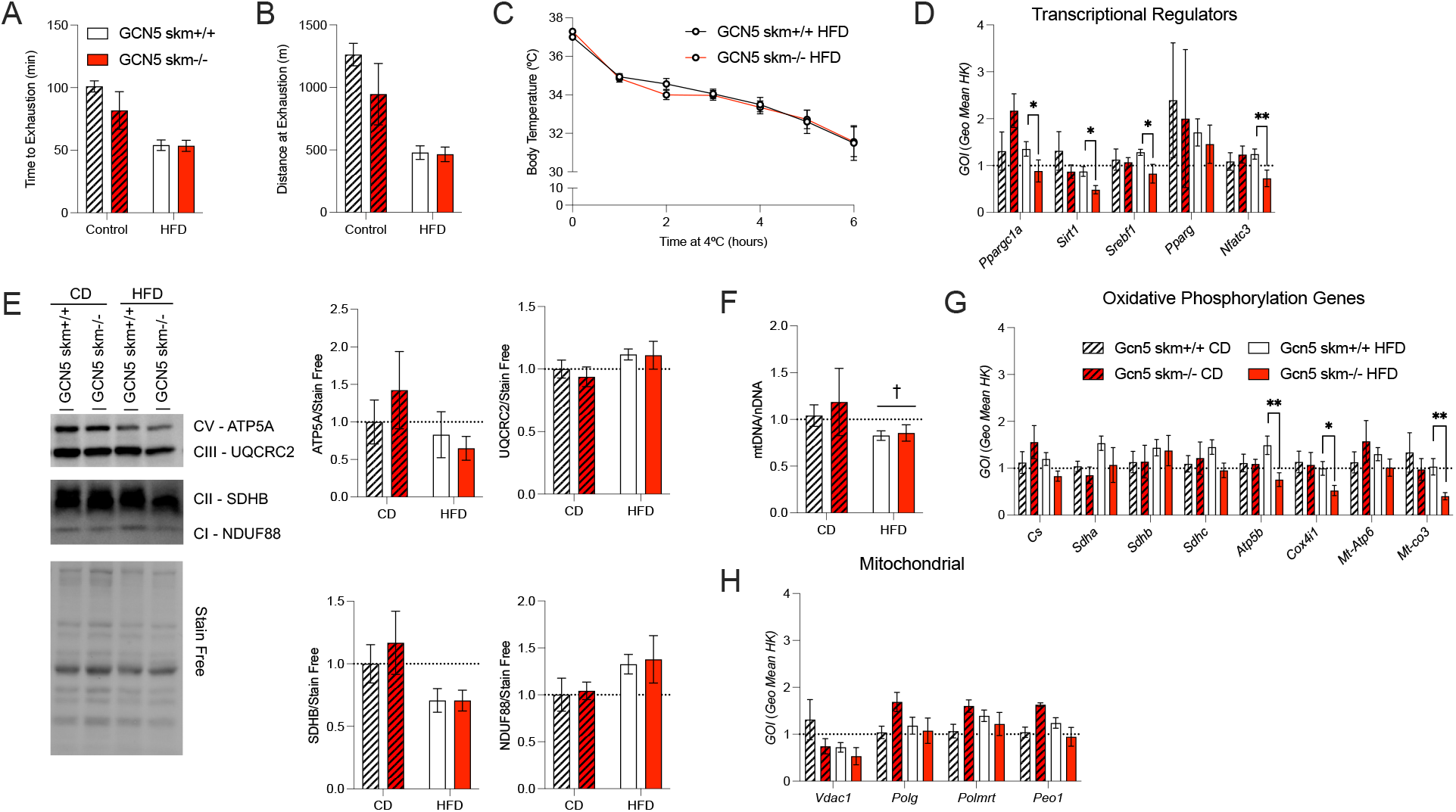
GCN5 ablated mice have similar exercise capacity, cold tolerance, and mitochondrial content to wildtype mice. There were no differences in time to exhaustion (A) or distance completed at exhaustion (B) between Gcn5 skm+/+ and Gcn5 skm-/- (n = 4-12). When acutely housed at 4°C, there were no differences in body temperature (C; n = 12). D) Quadriceps mRNA measurements of transcriptional regulators (n = 4-12). E) Representative western blot of mitochondrial oxidative phosphorylation proteins and quantification. F) Mitochondrial DNA levels in quadriceps muscle (n = 4-12). Quadriceps mRNA levels of oxidative phosphorylation genes (G) and mitochondrial genes (H; n = 4-12). Bars in graphs represent mean values and error bars are s.e.m. *: p < 0.05, ** p < 0.01, *** p < 0.001 and **** p < 0.0001 indicates significance by 1/2-way ANOVA and Bonferroni post-hoc tests.

As it has previously been shown that 1) both GCN5 and its homolog PCAF can regulate PGC-1α acetylation levels[5], 2) GCN5 levels increase in the absence of PCAF[17], and 3) other laboratories have suggested that other lysine acetyl transferases may be the major regulators of skeletal muscle acetylation levels[7], the effect of dual GCN5 and PCAF inhibition in skeletal muscle was examined. In support of PCAF compensating for the absence of GCN5, we observed a significant increase in *Kat2b* (gene for PCAF) mRNA levels in quadriceps muscle in control diet (CD)-fed mice (Fig. 4A). Interestingly, this compensation was not observed and was even reversed in HFD-fed mice (Fig. 4A).

**Figure 4.**
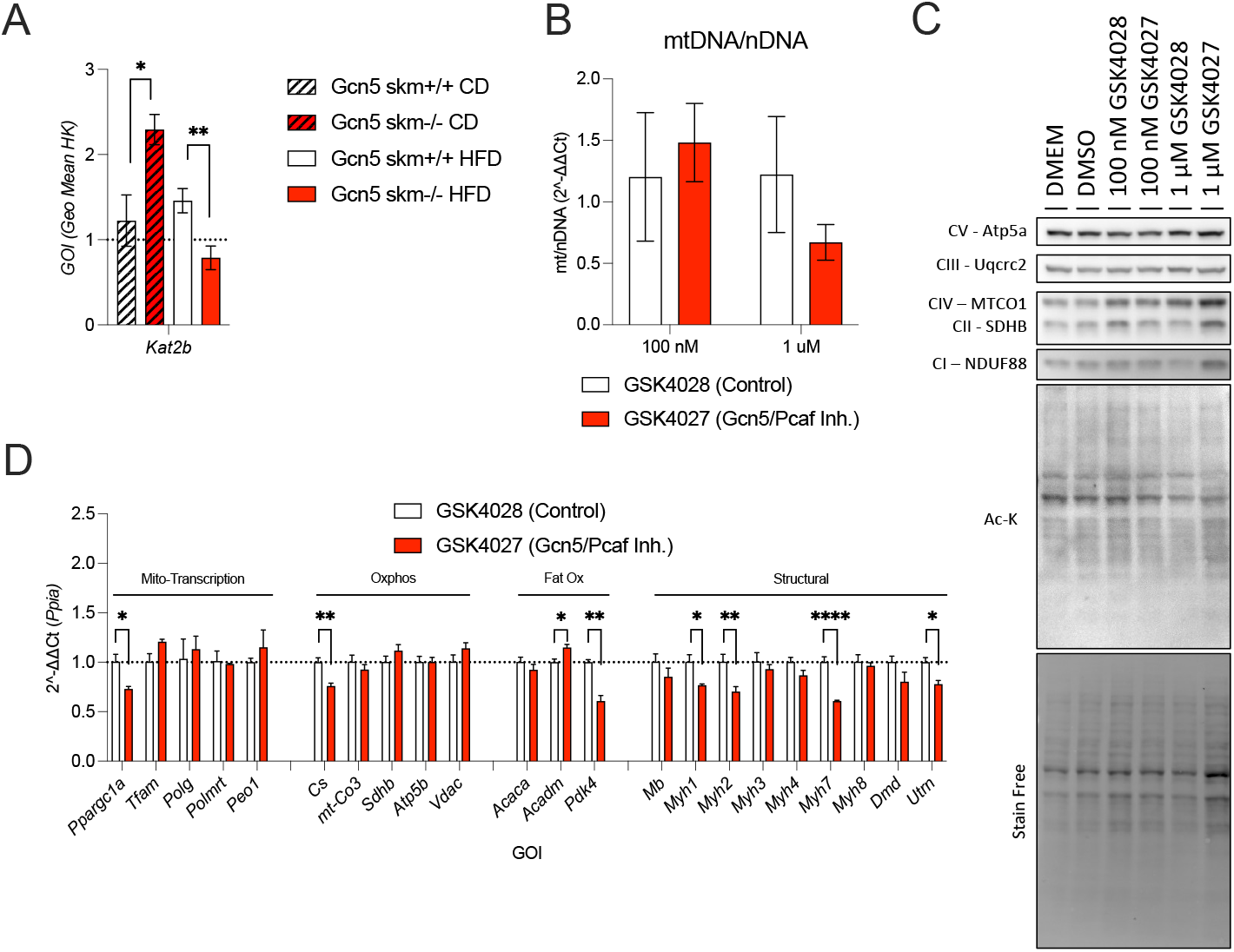
PCAF increases in response to Gcn5 ablation and inhibition of both Kat2a and Kat2b decreases the expression of many mitochondrial genes in myotubes. A) Kat2b levels are higher in Gcn5 skm-/- mice versus Gcn5 skm+/+ on a control diet but lower on the HFD (n = 4-12). GSK4027 (Gcn5/PCAF inhibitor) does not increase mitochondrial DNA (B) or oxidative phosphorylation protein expression (C), and decreases gene expression of mitochondrial gene transcription regulators, mitochondrial gene expression, fat oxidation genes and structural genes (E). Bars in graphs represent mean values and error bars are s.e.m. *: p < 0.05, ** p < 0.01, *** p < 0.001 and **** p < 0.0001 indicates significance by either student-s test, 1-, or 2-way ANOVA and Bonferroni post-hoc tests. Ac-K = Acetylated-Lysine

To test if inhibiting the acetylation-reading capabilities of GCN5 and PCAF resulted in changes in gene expression, GSK4027, a compound capable of inhibiting the homologous bromodomain of GCN5 and PCAF with similar affinity, and its inactive enantiomer, GSK4028[13], were applied to primary myotubes in culture for 72 hours. Subsequently, mitochondrial, and metabolic measures were assessed in these cells. Surprisingly, there were no differences in mitochondrial content, as measured by mtDNA levels (Fig. 4B) and oxidative phosphorylation protein/gene expression levels (Fig. 4C-D). Further and consistent with *Gcn5 skm-/-* mice (Fig. 3D), the mRNA levels of *Ppargc1a* and *Cs* decreased with GSK4027 treatment, but not that of *mt-Co3* or *Atp5b* (Fig. 4D). Similar to *Gcn5 skm-/-* mice and consistent with the observed decrease in fat oxidation in these mice, GSK4027 decreased expression of *Pdk4* (Fig.4D), although we did observe a small increase in the fatty acid oxidation gene *Acadm* (Fig.4D). Additionally, paralleling our published results in *Gcn5 skm-/-* mice, GSK4027 decreased expression of key structural and functional genes in skeletal muscle including numerous *Myh* genes and *Utrn* (Fig.4D). All told, GSK4027 inhibition of GCN5 and PCAF did not increase or decrease the expression of mitochondrial gene but did decrease skeletal muscle structural gene expression.

## 4 Discussion

### 4.1 Role of GCN5 in regulation macronutrient oxidation

HFD-feeding revealed a surprising decrease in fat oxidation and oxidative phosphorylation gene expression in *Gcn5 skm-/-* versus *Gcn5 skm+/+* mice. The observed decrease in fat oxidation, contrasts with gain of function models[16], where overexpression of GCN5 suppressed PGC-1a induced gene expression in C2C12-derived myotubes, and subsequently decreased fat oxidation rates. As GCN5 is normally present in the Spt-Ada-Gcn5 acetyltransferase (SAGA) complex, overexpression of GCN5 alone may produce GCN5 that is not associated with the SAGA complex, and consequently result in promiscuous acetylation of atypical target proteins by GCN5. This, however, would not be observed in loss of function models. Other GCN5 skeletal muscle knockout models or GCN5 knockout/Sirt1 overexpression models had no differences in metabolic substrate preferences with chow diets[6,7]. This would suggest that HFD-feeding activates novel roles for GCN5 and is consistent with previous observations that GCN5 levels increase with HFD-feeding. Together this would suggest that skeletal muscle GCN5 plays a larger role under HFD-feeding and may be required for full induction of fat oxidation.

Notably, gene expression of *Pdk4*, an important metabolic switch that inhibits pyruvate dehydrogenase, is decreased with *Gcn5* loss and dual GCN5 and PCAF inhibition, but only under HFD conditions or in cell culture conditions with high (25 mM) glucose levels. *Pdk4* KO mice have lower fat oxidation and increased glucose oxidation rates[18], thus presumably lower *Pdk4* levels would permit greater glucose oxidation. Interestingly, *Pdk4* is predicted to be a transcriptional target of YY1 (https://maayanlab.cloud/Harmonizome/gene/YY1)[19,20], which we and others have previously described to be a target of protein acetylation, and specifically by GCN5[10]. This would suggest that *Pdk4* expression may be regulated by GCN5, and the loss of *Gcn5* results in decreased *Pdk4* expression with a concurrent increase in glucose oxidation or decrease in fat oxidation.

### 4.2 Gcn5 loss does not promote mitochondrial gene expression even under HFD-feeding

Notably, our group and others have demonstrated that, in contrast with overexpression of GCN5, skeletal muscle loss of GCN5 has only minor effects on basal or exercise induced mitochondrial gene expression in chow fed animals [6,7,10]. Only combined *Sirt1* overexpression (~350 fold) and *Gcn5* gene ablation modestly increases mitochondrial content and fat oxidation rates [7].

As HFD-feeding increases mitochondrial content[21] and GCN5 levels[8], we expected that loss of *Gcn5* in skeletal muscle might synergistically enhance the HFD-induced increase in mitochondrial content; however, the opposite was observed. *Gcn5* loss and combined inhibition of GCN5 and PCAF in hyperglycemic primary cell culture decreased Sirt1, Ppargc1a and expression of many oxidative phosphorylation genes. Since combined over expression of *Sirt1* and loss of *Gcn5* is required to see an increase in mitochondrial content under chow-fed conditions[7], and we observed a decrease in *Sirt1* expression alone, this would suggest that *Gcn5* loss alone would not enhance mitochondrial content. Therefore, HFD with a loss of *Gcn5* resulted in a decrease in the expression of mitochondrial gene transcriptional regulators and subsequently mitochondrial gene expression.

### 4.3 Compensation by other Acetyltransferases

It has been suggested that other acetyltransferases, particularly PCAF, can compensate for the loss of GCN5 protein expression. This is supported by the high sequence homology (~73%) [11] between GCN5 and PCAF, as well as by observations that GCN5 increases in the absence of PCAF[17] and that loss of both PCAF and GCN5 is required to reduce acetylation of K9 on H3[22]. Consistent with this, we observed an increase in *Kat2b* mRNA levels in CD-fed *Gcn5 skm-/-* mice and no differences in overall acetylation levels. However, with HFD-feeding, there was no compensatory effect of *Kat2b* levels. Further, when inhibiting the bromodomain of both GCN5 and PCAF, we observed similar effects to the HFD-fed *in vivo* loss of *Gcn5* alone. This would suggest that PCAF compensation alone cannot explain the lack of mitochondrial gene expression enhancement, consistent with the requirement for *Sirt1* overexpression. This, however, does not eliminate the possibility that other acetyltransferases may play a larger role in regulating the activity of PGC-1a or other factors.

## 5 Conclusions

HFD-feeding revealed novel roles for GCN5 in skeletal muscle. GCN5 loss reduces expression of *Pdk4* expression and increases glucose oxidation with HFD-feeding. Further, dual inhibition of GCN5 and PCAF did not increase mitochondrial content but did decrease genes inhibiting glucose oxidation. Therefore, GCN5 and PCAF do not appear to be major inhibitors of mitochondrial content but do appear to regulate macronutrient oxidation preference.

## 6 Acknowledgements

The authors would like to thank the staff of the University of Ottawa Animal Behaviour and Physiology Core and the Louis Pelletier Histology Core. A.E. Green is the recipient of a University of Ottawa Brain and Mind Research Institute uOttawa Eric Poulin Centre for Neuromuscular Disease Scholarship in Translational Research Award. K.J. Menzies received funding from the Canadian Institutes of Health Research (MOP 159455) and the Natural Sciences and Engineering Research Council of Canada Collaborative Research and Training Experience program (Metabolomics Advanced Training and International Exchange program) and discovery grants (RGPIN 2018-06838, DGECR 2018-00012). H. Zhang was funded by the National Natural Science Foundation of China (31871370 and 32000840), the Natural Science Foundation of Guangdong Province (2018A030313655 and 2019A1515011342), Science and Technology Program of Guangzhou (202002030429). D. Ryu was supported by the Basic Science Research Program of the Korean government (Ministry of Science and ICT grants NRF-2020R1A2C2010964 and 2021R1A5A8029876). J. Auwerx received funding from the École Polytechnique Fédérale de Lausanne, the European Research Council (ERC-AdG-787702), the Swiss National Science Foundation (SNSF 31003A_179435), and the Global Research Laboratory grant of the National Research Foundation of Korea (NRF 2017K1A1A2013124)

The authors declare no competing financial interests related to this work.

## 8 Appendix A: Supporting Figures

**Figure S 1.**
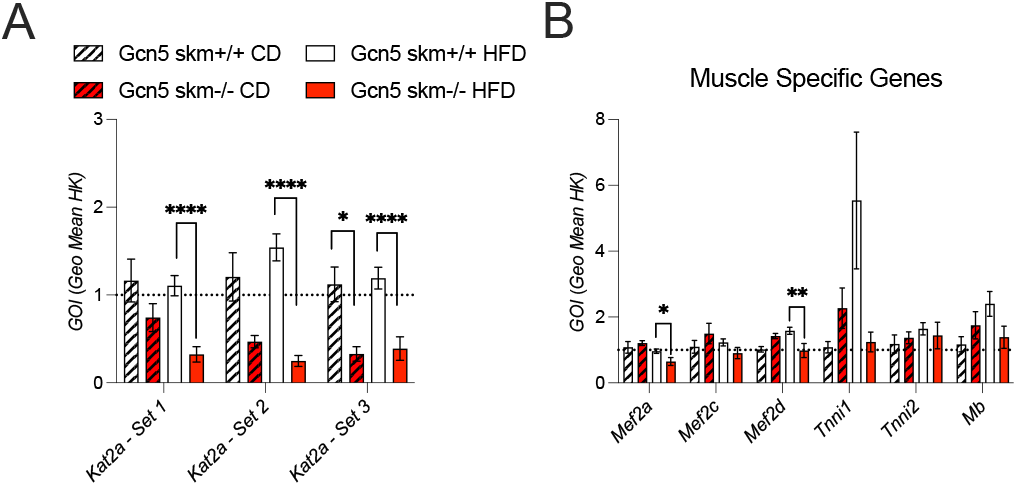
A) Expression of Kat2a (Gcn5) levels using 3 different primer sets and muscle specific genes (B). Bars in graphs represent mean values and error bars are s.e.m. *: p < 0.05, ** p < 0.01, *** p < 0.001 and **** p < 0.0001 indicates significance by either 1-, or 2-way ANOVA and Bonferroni post-hoc tests.

## 9 Appendix A: Supporting Tables

**Table S 1.**
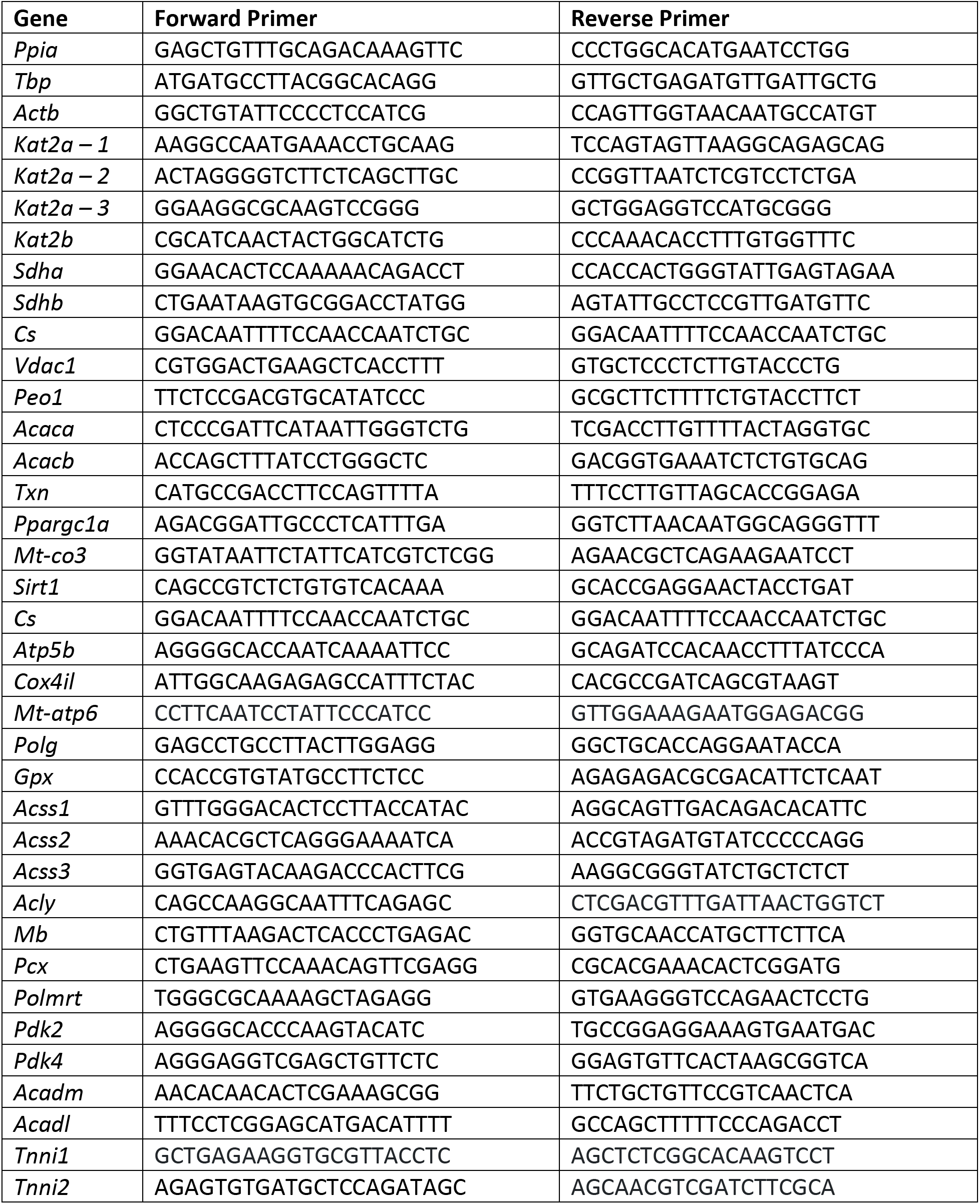
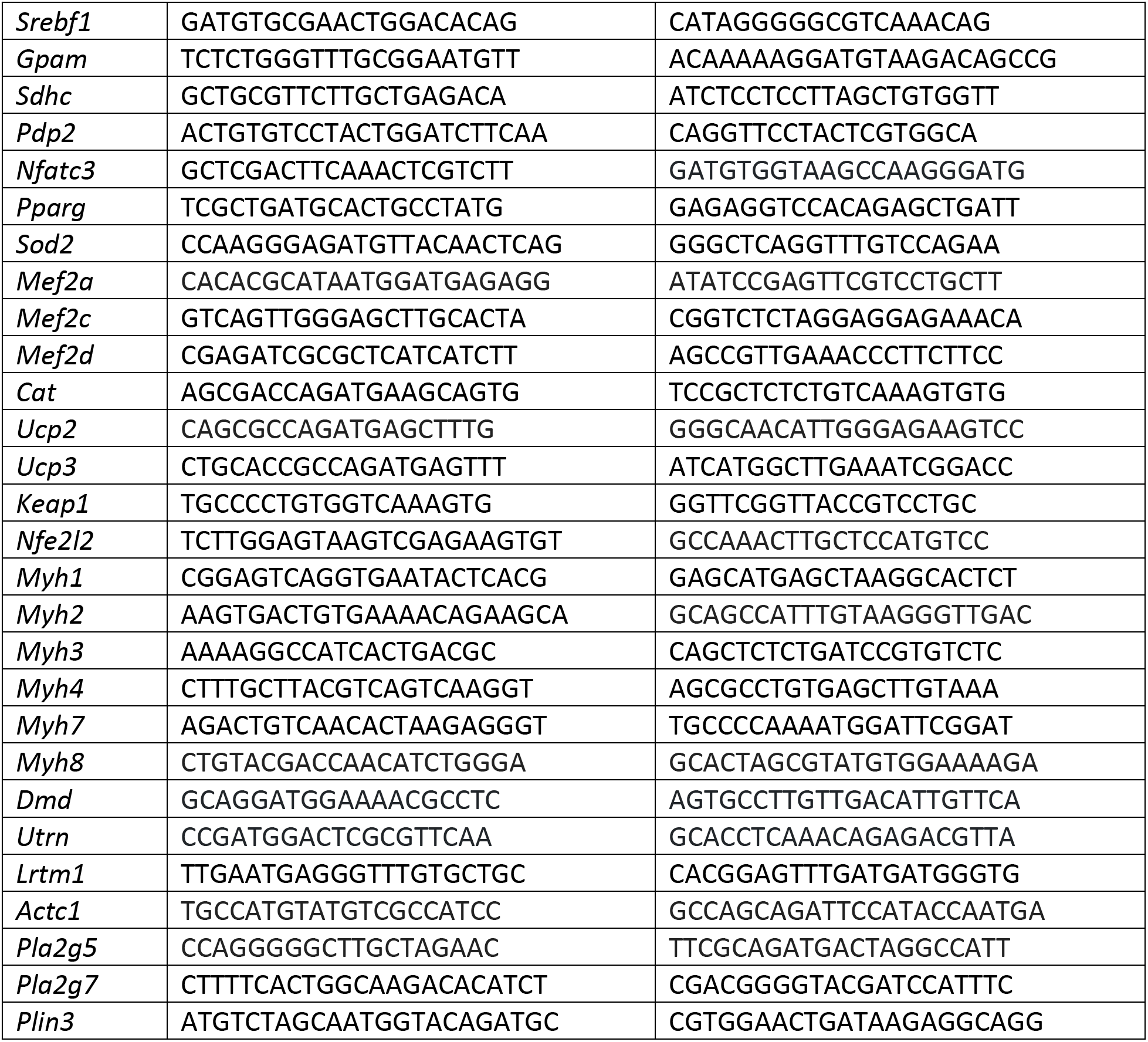
Primer sequences for RNA quantification using SYBR-Green Technology.

**Table S 2.** Taqman primers used for RNA quantification.

| Gene | Taqman Probe ID |
| --- | --- |
| *Kat2a* | Mm02342430_g1 |
| *Ppia* | Mm02342430_g1 |

**Table S 3.**
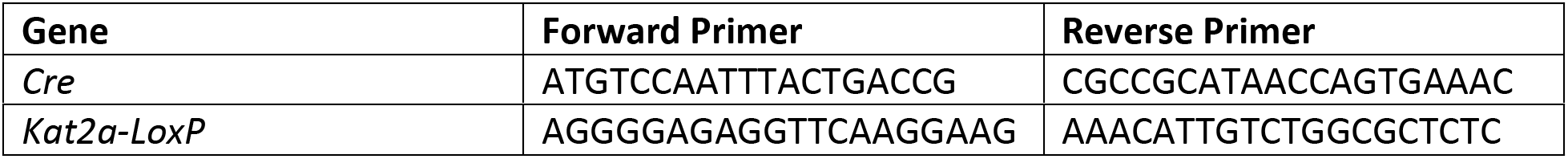
Primer sequences used for genotyping.

**Table S 4.**
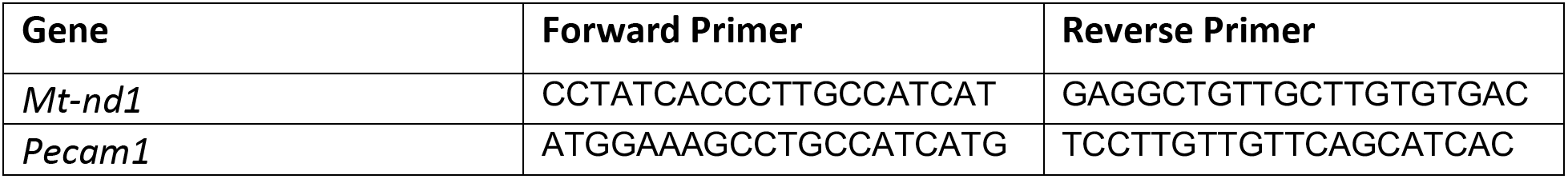
Primer sequences used for mtDNA to nDNA ratio quantification.

**Table S 5.**
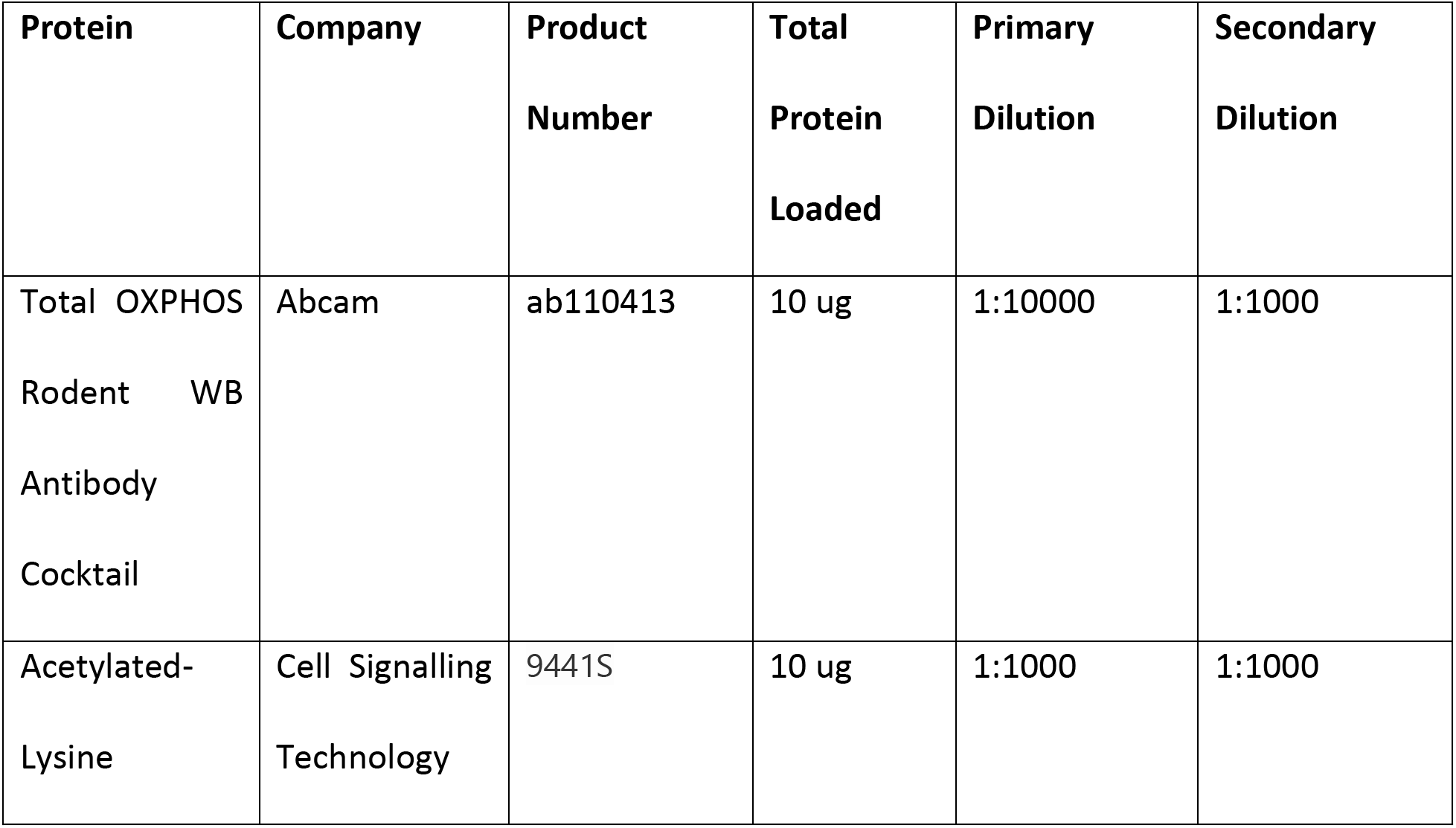
List of antibodies and dilutions used for protein quantification.

## Notes

### Competing Interest Statement

The authors have declared no competing interest.

